# Alterations in the preneoplastic breast microenvironment of *BRCA1/2* mutation carriers revealed by spatial transcriptomics

**DOI:** 10.1101/2023.05.24.542078

**Authors:** Anthony Caputo, Kavya Vipparthi, Peter Bazeley, Erinn Downs-Kelly, Patrick McIntire, Ying Ni, Bo Hu, Ruth A. Keri, Mihriban Karaayvaz

## Abstract

Breast cancer is the most common cancer in females, affecting one in every eight women and accounting for the majority of cancer-related deaths in women worldwide. Germline mutations in the *BRCA1* and *BRCA2* genes are significant risk factors for specific subtypes of breast cancer. *BRCA1* mutations are associated with basal-like breast cancers, whereas *BRCA2* mutations are associated with luminal-like disease. There are currently few chemoprevention strategies available for *BRCA1*/*2* mutation carriers, and irreversible prophylactic mastectomy is the primary option. Designing chemo-preventive strategies requires an in-depth understanding of the physiological processes underlying tumor initiation. Here, we employ spatial transcriptomics to investigate defects in mammary epithelial cell differentiation accompanied by distinct microenvironmental alterations in preneoplastic breast tissues from *BRCA1*/*2* mutation carriers and normal breast tissues from non-carrier controls. We uncovered spatially defined receptor-ligand interactions in these tissues for the investigation of autocrine and paracrine signaling. We discovered that β1-integrin-mediated autocrine signaling in *BRCA2*-deficient mammary epithelial cells differs from *BRCA1*-deficient mammary epithelial cells. In addition, we found that the epithelial-to-stromal paracrine signaling in the breast tissues of *BRCA1*/*2* mutation carriers is greater than in control tissues. More integrin-ligand pairs were differentially correlated in *BRCA1*/*2*-mutant breast tissues than non-carrier breast tissues with more integrin receptor-expressing stromal cells. These results reveal alterations in the communication between mammary epithelial cells and the microenvironment in *BRCA1* and *BRCA2* mutation carriers, laying the foundation for designing innovative breast cancer chemo-prevention strategies for high-risk patients.

## Introduction

Breast cancer is the most common cancer in females, affecting one in every eight women and accounting for the majority of cancer-related deaths in women worldwide [1]. Germline predisposition, reproductive history, and mammographic density are the best-known predictors of breast cancer risk. A single full-term pregnancy in early adulthood reduces the lifetime risk of hormone receptor-positive postmenopausal breast cancer by approximately twofold [2]. High mammographic density is a strong predictor of breast cancer risk. A major group of women who are at high risk for breast cancer are those with germline mutations in the *BRCA1* or *BRCA2* genes. Cumulative breast cancer risk is 72% for *BRCA1* mutation carriers and 69% for *BRCA2* mutation carriers by the age of 80 [3]. There are currently few chemoprevention strategies available for *BRCA1*/*2* mutation carriers, and irreversible prophylactic mastectomy is the primary option [4]. Although prophylactic mastectomy is a very effective preventative measure, many women are reluctant to undergo such invasive surgery. Thus, there is a clear need to discover novel chemo-preventive strategies for these high-risk women. Importantly, breast cancers arising in *BRCA1* and *BRCA2* mutation carriers differ in their cancer subtype classification. Germline mutations in *BRCA1* are associated with hormone-receptor negative, basal-like breast cancers, whereas those in *BRCA2* are associated with hormone receptor-positive, luminal-like breast cancers [5-8]. Basal-like breast cancers are proliferative, high-grade and associated with a poor prognosis. In these cancers, therapeutic options include chemotherapy, PARP inhibitors and immunotherapy [9, 10]. On the other hand, luminal-like breast cancers are often low-grade, differentiated tumors expressing hormone receptors. They can be treated with hormonal therapy and generally display a relatively favorable survival [11, 12]. Therefore, the etiology and treatment of breast cancer varies significantly between *BRCA1* and *BRCA2* germline mutation carriers. However, the underlying biological mechanism remains largely unknown.

The human mammary gland comprises of epithelial and stromal cells communicating with each other via extracellular matrix (ECM) [13]. The mammary epithelium consists of two differentiated cell types organized into two cell layers, an inner layer of luminal epithelial cells and an outer layer of basal (myoepithelial) cells in direct contact with the basement membrane [14]. Prior transcriptomic analyses of patient specimens have revealed loss of lineage fidelity in germline *BRCA1/2* mutation carriers’ preneoplastic mammary epithelial cells [15-19]. However, cell fate decisions are driven by changes in environmental signals [20]. The role of the microenvironment in mammary epithelial cell fate determination in *BRCA1/2* mutation carriers has not been investigated. Here, we used NanoString GeoMx Digital Spatial Profiling (DSP) technique to allow spatial characterization of the epithelia and microenvironment in normal and preneoplastic *BRCA1/2*-mutant (*BRCA1/2*^*mut/+*^) human breast tissues. We discovered spatially defined receptor-ligand interactions for autocrine and paracrine signaling, enabling comprehensive understanding of communication processes between mammary cells. We discovered that while *BRCA2* deficiency activates *β*1-integrin mediated mechano-transduction in preneoplastic human mammary epithelial cells, *BRCA1* deficiency does not, implying that *BRCA1*- and *BRCA2*-deficient cells communicate with the ECM in fundamentally different ways. On the other hand, *BRCA1*- and *BRCA2*-deficient preneoplastic human breast tissues are comparable in terms of paracrine signaling between epithelial cells and surrounding stromal cells, with *BRCA1/2*^*mut/+*^ breast tissues having higher paracrine signaling than control tissues. These findings pave the way for developing novel breast cancer chemoprevention strategies in high-risk patients.

## Results

### Digital spatial transcriptomic profiling of human breast tissues

The study cohort included 12 patients (Fig. 1a and Supplementary Table 1). Noncancerous human breast tissues were collected from germline *BRCA1/2* mutation carriers who underwent prophylactic mastectomy and from non-mutation carriers who underwent elective breast reduction mammoplasty (wild type, WT). Since genetic breast cancers have a younger age onset, we collected samples from patients who were younger than 50 years of age. Three formalin-fixed paraffin-embedded (FFPE) tissue sections (each from one wild-type, one *BRCA1* and one *BRCA2* mutation carrier patient) were placed on one slide for performing RNA sequencing using the NanoString GeoMx DSP platform. After quality control filtering, expression data for 142 regions-of-interest (ROIs) across 11,833 genes were selected for downstream analysis (Supplementary Fig. 1a). The raw sequencing counts were normalized using the third quartile of expression (Q3 normalization) and log2-transformed for all subsequent analyses (Supplementary Fig. 1b, c). Principal component analysis (PCA) showed batch effects which were corrected for in the downstream analyses (Supplementary Fig 1d). Key findings were validated experimentally to ensure quality.

**Fig. 1:**
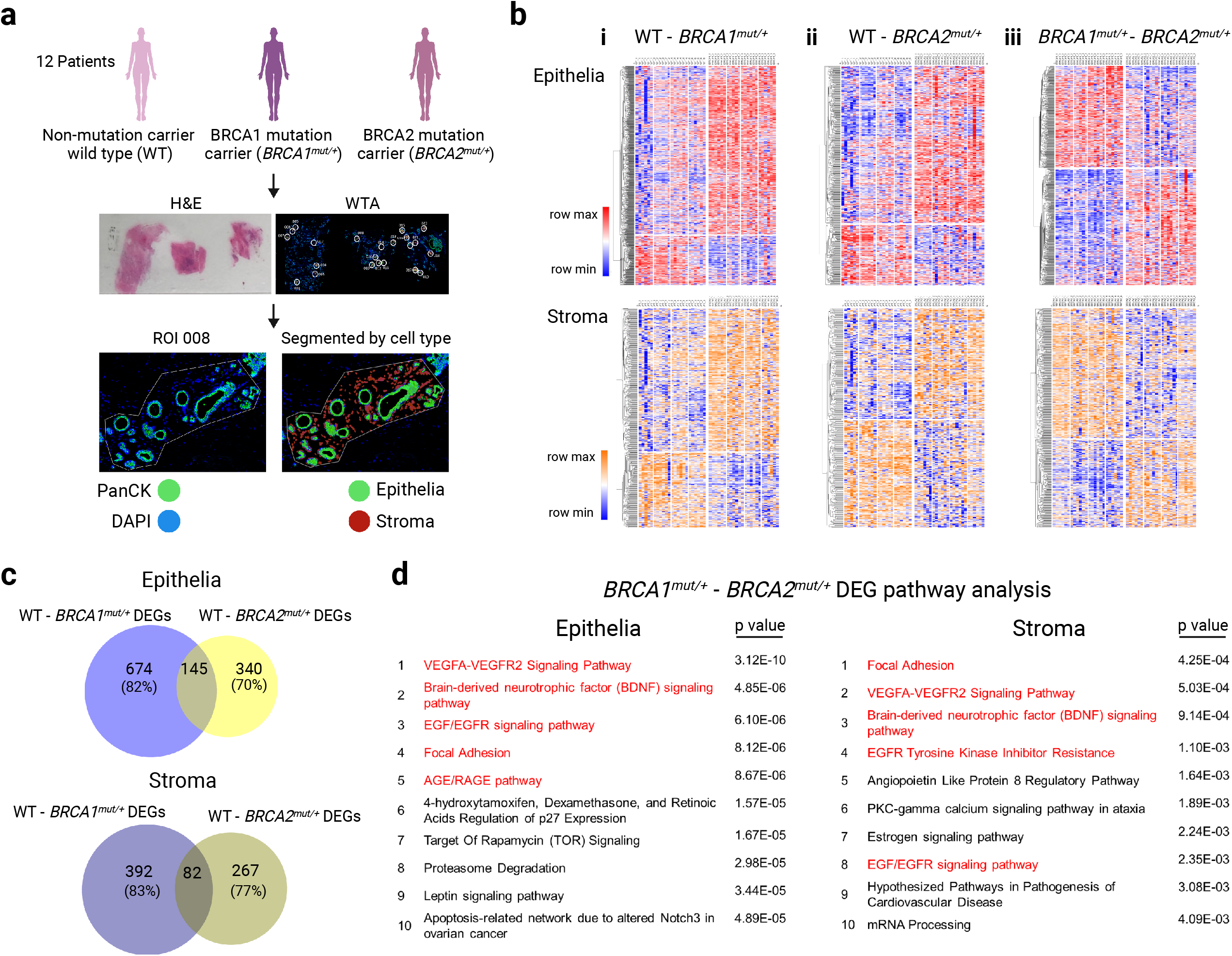
Digital spatial profiling of human breast tissues. **a** Schematic showing the study’s design and workflow. Human mammary epithelial and stromal cells from *BRCA1/2* mutation carrier prophylactic mastectomy and control (wild type (*WT*)) elective mammoplasty cases were used for digital spatial profiling (DSP) with the Nanostring GeoMx human whole-transcriptome atlas (WTA; 18,815 genes). Representative hematoxylin and eosin (H&E) stained tissue section and immunofluorescence image (WTA DSP) of consecutive sections from the same FFPE block are shown. Selected regions of interest (ROIs) are circled. Blue, SYTO13 (nuclear stain); green, anti-panCK (epithelial marker). Example ROI with segmentation masks used to enrich for epithelial (panCK positive and nuclear stain positive) and stromal (panCK negative and nuclear stain positive) cells. **b** Heatmaps created with Morpheus (https://software.broadinstitute.org/morpheus/) using a relative scale color scheme to show differentially expressed genes (DEGs) between genotypes (i: *WT*-*BRCA1*^*mut/+*^; ii: *WT*-*BRCA2*^*mut/+*^; iii: *BRCA1*^*mut/+*^-*BRCA2*^*mut/+*^) with nominal p values < 0.05 in the epithelia (top) and stroma (bottom). A total of 142 ROIs were analyzed. Each column represents either an epithelia or stroma segment. **c** Venn diagrams contrasting DEGs from the *WT*-*BRCA1*^*mut/+*^ (i) and *WT*-*BRCA2*^*mut/+*^ (ii) comparisons from ‘b’. Number of genes common and unique to each comparison are shown within circles, along with percentages of genes unique to each comparison. **d** Top-ranked pathways from WIKI pathway analysis of *BRCA1*^*mut/+*^-*BRCA2*^*mut/+*^ DEGs (iii) in epithelia (left) and stroma (right) from ‘b’. Pathways for growth factor signaling and focal adhesion are highlighted in red. WT: n = 4 patients; *BRCA1*^*mut/+*^: n= 4 patients; *BRCA2*^*mut/+*^: n = 4 patients.

To decipher expression programs in spatially organized epithelial and surrounding stromal cells, histological analyses following hematoxylin and eosin staining and immunostaining with the epithelial cell marker pan-cytokeratin (PanCK) and SYTO13 to mark the cell nuclei were used. Cell-type segments were created based on PanCK staining (epithelia segment is positive for PanCK and SYTO13, while stroma segment is negative for PanCK and positive for SYTO13) (Supplementary Fig. 1e). Our sequencing data contained 40% epithelial cells and 60% of stromal cells, according to segmentation masks that were utilized to aggregate the percent of total segment area occupied by each compartment across all ROIs (Supplementary Fig. 1f). We identified more nominally differentially expressed genes (DEGs) in epithelial cells compared to stromal cells in pairwise comparisons across genotypes (Fig. 1b). A total of 819 genes were found to be nominally differentially expressed in epithelial cells between the WT and *BRCA1*^*mut/+*^ groups, 485 genes between the WT and *BRCA2*^*mut/+*^ groups and 652 genes between the *BRCA1*^*mut/+*^ and *BRCA2*^*mut/+*^ groups (unadjusted p < 0.05). A total of 474 genes were found to be nominally differentially expressed in stromal cells between WT and *BRCA1*^*mut/+*^ groups, 349 genes between the WT and *BRCA2*^*mut/+*^ groups and 494 genes between the *BRCA1*^*mut/+*^ and *BRCA2*^*mut/+*^ groups (unadjusted p < 0.05). Intriguingly, DEGs identified in WT-*BRCA1*^*mut/+*^ comparison did not overlap with those identified in the WT-*BRCA2*^*mut/+*^ comparison (∼80% genes were unique), suggesting that the expression patterns of preneoplastic mammary cells from *BRCA1* and *BRCA2* mutation carriers are significantly different. Pathway analyses of DEGs in *BRCA1*^*mut/+*^-*BRCA2*^*mut/+*^ groups showed that integrin and growth factor signaling is differentially regulated between *BRCA1* and *BRCA2* mutation carrier patients (Fig. 1d and Supplementary Fig. 1g).

### Autocrine signaling pathways distinguish between *BRCA1*- and *BRCA2*-deficient preneoplastic human mammary cells

To uncover autocrine signaling in epithelia and stroma compartments of *BRCA1/2*-deficient human breast tissues, we identified spatially defined receptor-ligand pairs co-expressed in either epithelial or stromal cells [21]. Integrins constituted the majority of the receptor family that was found to be differentially correlated in genotype comparisons (Fig. 2a). *β*1-integrin (*ITGB1*) represents the predominantly expressed integrin in mammary cells and it is crucial for mammary gland development and differentiation [22, 23]. *β*1-integrin expression was reduced in *BRCA1*^*mut/+*^ epithelial and stromal cells when compared with control and *BRCA2*^*mut/+*^ cells (Supplementary Fig. 2a). Therefore, we explored *β*1-integrin-ligand pairs in control and *BRCA1*/*2-* deficient human breast epithelial and stromal cells. *β*1-integrin-ligand pairs were more prevalent in mammary epithelial cells than in stromal cells, and they were the most differentially correlated in the *BRCA1*^*mut/+*^-*BRCA2*^*mut/+*^ comparison (Fig. 2b). *β*1-integrin expression correlated with the expression of matrix degrading enzymes (*ADAM17*), angiogenesis genes (*THBS1*) or growth factors/chemokines (*CSF2, CXCL12*) in *BRCA1*^*mut/+*^ cells. On the other hand, *β*1-integrin expression correlated with the expression of type I collagen (*COL1A1*) and laminin genes (*LAMA4, NID1*) in *BRCA2*^*mut/+*^ cells. Integrins are heterodimeric transmembrane receptors made up of αand *β*subunits [24]. *β*1 integrin becomes a collagen receptor when heterodimerizes with the α11subunit [25]. Expression of α11subunit gene *ITGA11* also correlated with the expression of type 1 collagen genes (*COL1A1* and *COL1A2*) in *BRCA2*^*mut/+*^ mammary cells.

**Fig. 2:**
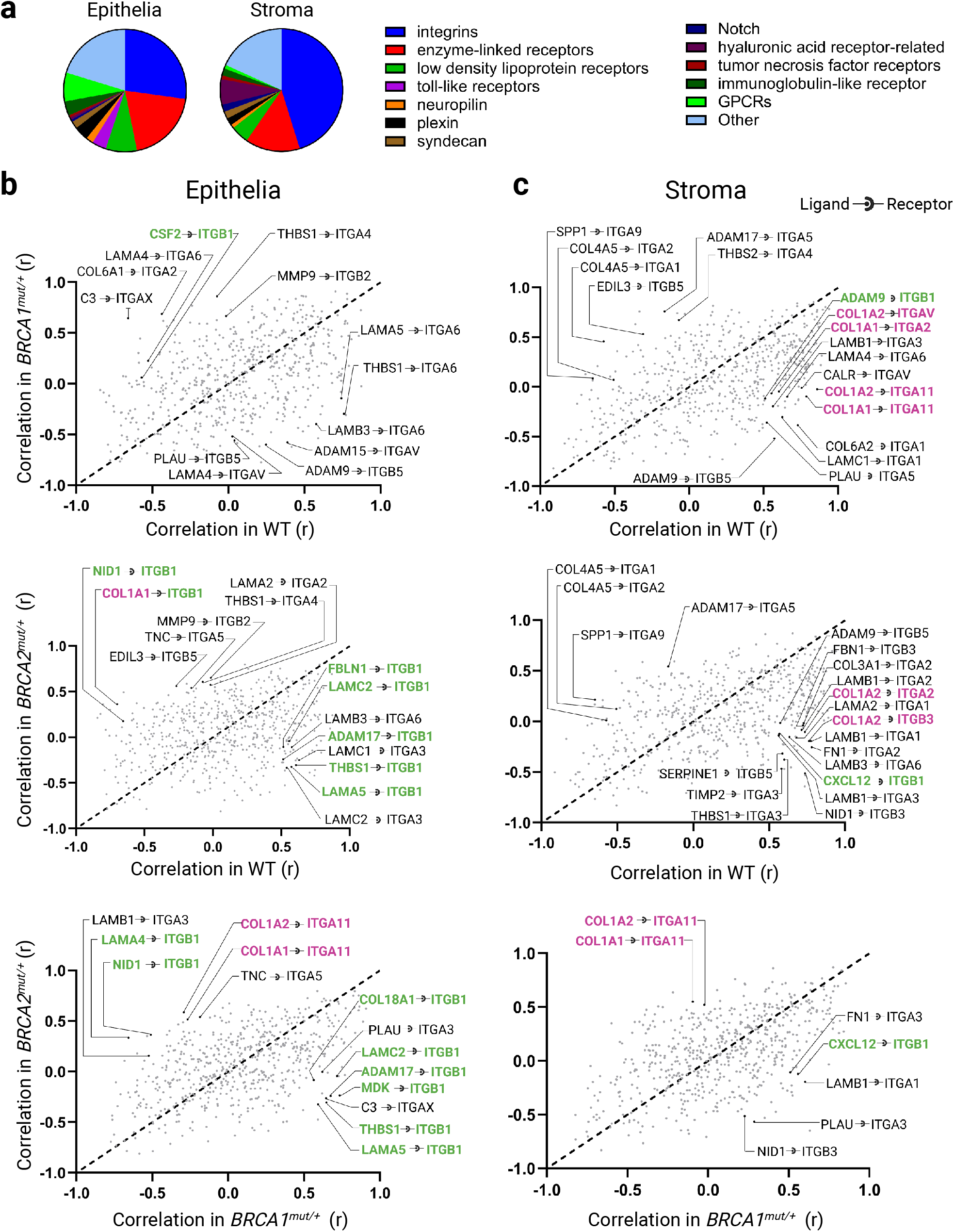
Spatial analysis within epithelia and stroma compartments reveals altered receptor-ligand interactions with the ECM in *BRCA1* and *BRCA2* mutation carriers. **a** Based on the receptor expression, pie charts illustrate the prevalence of spatially correlated receptor-ligand pairs within epithelia and stroma compartments. Pearson correlation coefficient (r) of expression of receptor-ligand pairs within **(b)** epithelia and **(c)** stroma compartments are represented with gray dots. ITGB1-cognate ligand pairs that were differentially correlated in pairwise comparison across genotypes are labeled in green. Collagen type I-cognate receptor pairs that were differentially correlated in pairwise comparison across genotypes are labeled in pink.

Enzyme-linked receptors constituted the second most abundant receptor family that was found to be differentially correlated in genotype comparisons (Supplementary Fig. 2b). TGF*β*pathway receptors were most significantly altered among the enzyme-linked receptors in genotype comparisons. Importantly, *TGFBR3*-*TGFB2, BMPR2*-*BMP2* and *BMPR2*-*RGMB* pairs were shown to be differentially correlated in *BRCA2*^*mut/+*^ mammary epithelial cells compared to *BRCA1*^*mut/+*^ cells. TGF-*β*signaling is a potent inducer of collagen production by both epithelial and stromal cells in culture [26]. Integrin binding to collagen generates a mechanical force in the ECM which is linked to the cytoskeletal tension and activates mechano-signaling pathways [27, 28]. Interestingly, several actin/myosin genes were up-regulated in *BRCA2*^*mut/+*^ mammary epithelial cells compared to *BRCA1*^*mut/+*^ epithelial cells (Supplementary Fig. 2c). On the other hand, expression of the matrix metalloproteinases MMP3 and MMP8, which are collagen degrading enzymes, were increased in *BRCA1*^*mut/+*^ epithelial cells (Supplementary Fig. 2a) These findings suggest that ECM remodeling mechanisms differ significantly between *BRCA1*^*mut/+*^ and *BRCA2*^*mut/+*^ preneoplastic human mammary cells, with *BRCA1*^*mut/+*^ cells favoring ECM breakdown and *BRCA2*^*mut/+*^ cells favoring force-mediated ECM modification.

### *β*1 integrin-mediated mechano-signaling is diminished in *BRCA1*^*mut/+*^ preneoplastic mammary epithelial cells compared to *BRCA2*^*mut/+*^ preneoplastic mammary epithelial cells

Cells can detect and interpret the biological information in the ECM by adhering to it via integrin-based adhesions and eliciting a series of dynamic signaling events known as mechano-signaling. Integrins are transmembrane ECM receptors that function as mechano-transducers and integrin clustering activates growth factor-ERK-dependent cell growth [29, 30]. In this context, we asked whether *β*1 integrin-mediated mechano-signaling was differentially regulated in response to *BRCA1* or *BRCA2* loss. Our data showed that expression of activated *β*1 integrin and phosphorylated ERK was decreased with *BRCA1* loss-of-function compared to *BRCA2* loss-of-function in primary human mammary epithelial cells (Fig. 3 and Supplementary Fig. 3).

**Fig. 3:**
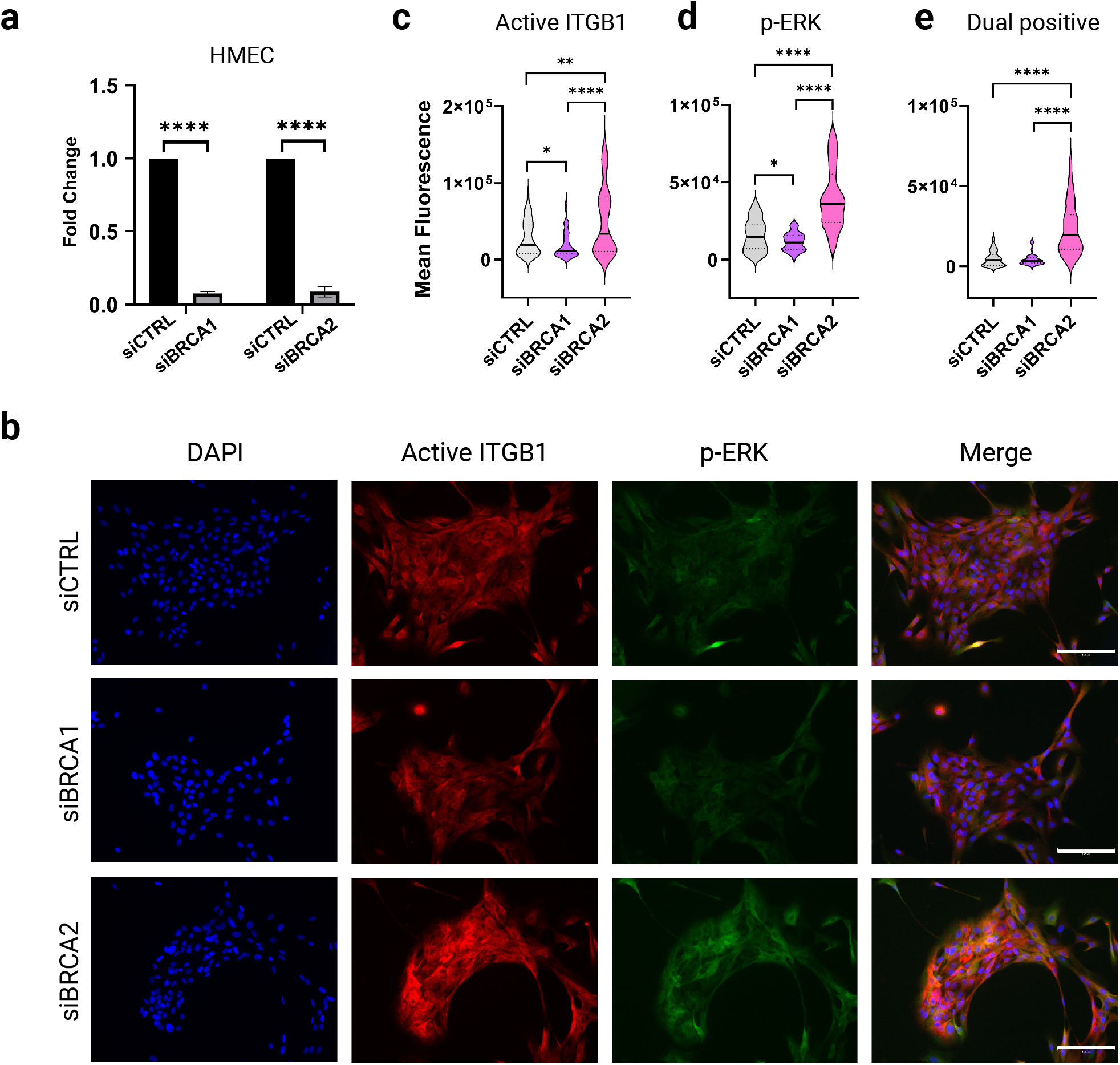
*BRCA2* deficiency increases *β*1 integrin-mediated mechano-signaling compared to *BRCA1* deficiency. **a** qRT-PCR analysis shows siRNA mediated knockdown of *BRCA1* and *BRCA2* genes in HMEC cells. p-values by unpaired t-test. * p < 0.05, ** p < 0.01 and **** p < 0.0001. **b** Representative images from immunofluorescence staining with an antibody to β1-integrin (active conformation) and phosphorylated ERK 1/2 (p-ERK) of HMEC cells following BRCA1 and BRCA2 knockdown. Scale bar = 150 μm. Violin plots show the quantification of staining intensity for **(c)** active β1-integrin, **(d)** p-ERK and **(e)** co-stained β1-integrin and p-ERK. Experiments were repeated two times independently. In each experiment, at least 15-20 images per condition were analyzed.

### Paracrine signaling is enhanced in both *BRCA1*^*mut /+*^and *BRCA2*^*mut/+*^ human breast tissues

Paracrine signaling mechanisms are essential for both normal mammary gland development and breast cancer [31]. To reveal the role of paracrine signaling in mammary epithelial cell fate determination in *BRCA1/2*^*mut/+*^ human breast tissues, we identified spatially defined receptor-ligand pairs co-expressed across ROIs in epithelial and stromal cells. In other words, we assessed the expression of a receptor in one compartment (epithelium or stroma) and its cognate ligand expression in the other compartment, or vice versa. Similar to autocrine signaling, integrins and enzyme-linked receptors comprised the majority of the receptor family in paracrine signaling that were found to be differentially correlated in pairwise genotype comparisons. As opposed to autocrine signaling, syndecans were a significant receptor family in paracrine signaling between epithelial and stromal cells (compare Fig. 4a and Fig. 2a). We discovered that *BRCA1/2*^*mut/+*^ human breast tissues exhibit greater integrin-dependent paracrine signaling than control tissues (Fig. 4b and 4c). Interestingly, in *BRCA1/2*^*mut/+*^ breast tissues, integrin and syndecans function more as stromal receptors than epithelial receptors, in striking contrast to control tissues. The *ITGA6*-*COL6A1* was one such receptor-ligand pair that was found to be differentially correlated between control and *BRCA1/2*^*mut/+*^ human breast tissues. In control tissues, epithelial *ITGA6* expression was positively correlated with stromal *COL6A1* expression; however, in *BRCA1/2*^*mut/+*^ breast tissues, this correlation disappeared. Consistently, stromal *ITGA6* expression was negatively correlated with epithelial *COL6A1* expression in control tissues; however, this correlation was absent in *BRCA1/2*^*mut/+*^ breast tissues (Supplementary Fig. 4a). We quantified ITGA6 immunostaining in mammary epithelial cells and type VI collagen immunostaining in the adjacent stroma using indirect immunofluorescence analysis, allowing us to validate our spatial transcriptomics findings (Fig. 5a) at the protein level (Fig 5b and 5c). Although comparable results were observed for enzyme-linked receptors, fewer receptor-ligand pairs with differential correlations were identified compared to integrin receptors in *BRCA1/2*^*mut/+*^ breast tissues (Supplementary Fig. 4b). *FGFR2*-*FGF2* was one such receptor-ligand pair that functioned more as stromal receptors than epithelial receptors in *BRCA1/2*^*mut/+*^breast tissues. Similar correlation patterns were observed between the expression of *FGFR2*-*FGF2* receptor-ligand and *ITGA6*-*COL6A1* (Supplementary Fig. 4c). Together, these data indicate that stromal cells in *BRCA1*/*2*^*mut/+*^ human breast tissues may have more active signaling with the microenvironment compared to stromal cells in control human breast tissues.

**Fig. 4:**
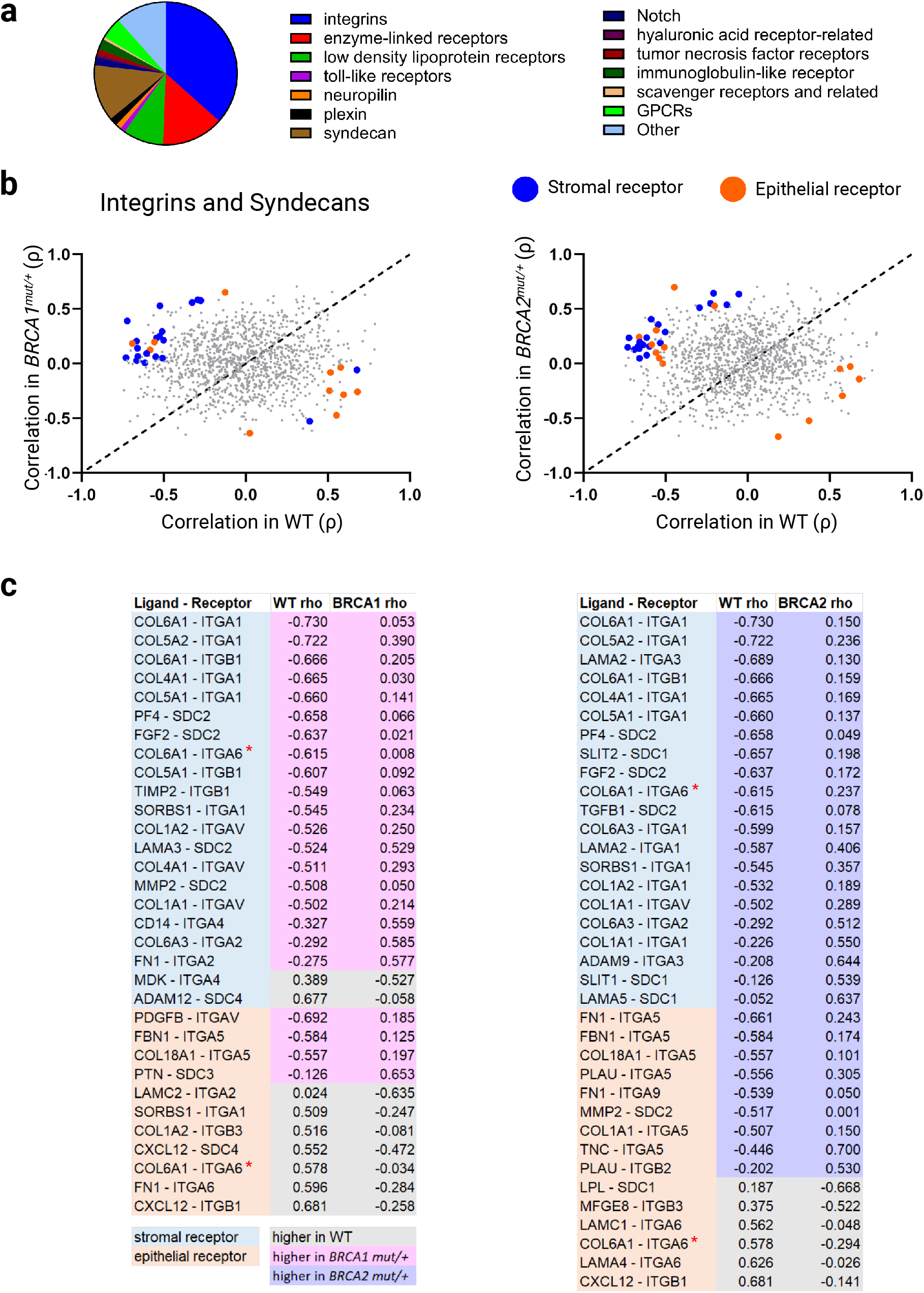
Spatial analysis across epithelia and stroma compartments reveals increased signaling of stromal cells with the ECM in *BRCA1*/*2* mutation carriers. **a** Based on the receptor expression, pie charts illustrate the prevalence of spatially correlated receptor-ligand pairs across epithelia and stroma compartments. **b** Spearman’s rho of expression of receptor-ligand pairs between epithelia and stroma compartments of WT, *BRCA1*^*mut/+*^ and *BRCA2*^*mut/+*^ patients are represented with gray dots. Differentially correlated integrin or syndecan-ligand pairs in pairwise comparisons of WT-*BRCA1* and WT-*BRCA2* are represented as colored dots; blue indicates a stromal receptor and orange indicates an epithelial receptor. **c** List of integrin or syndecan-ligand pairs from ‘b’. Background color of the receptor-ligand pairs: blue indicates higher stromal receptor expression and orange indicates higher expression of epithelial receptors. Background color of the correlation values: pink indicates correlation is higher in *BRCA1*^mut/+^ tissues, purple indicates correlation is higher in *BRCA2*^*mut/+*^ tissues, and gray indicates correlation is higher in WT tissues. Notably, the majority of blue receptor-ligand pairs have pink or purple correlation values, denoting increased signaling of stromal cells with the ECM through expression of integrins and syndecans. The red stars show the ITGA6-COL6A1 receptor-ligand pair that was investigated further in Fig. 5.

**Fig. 5:**
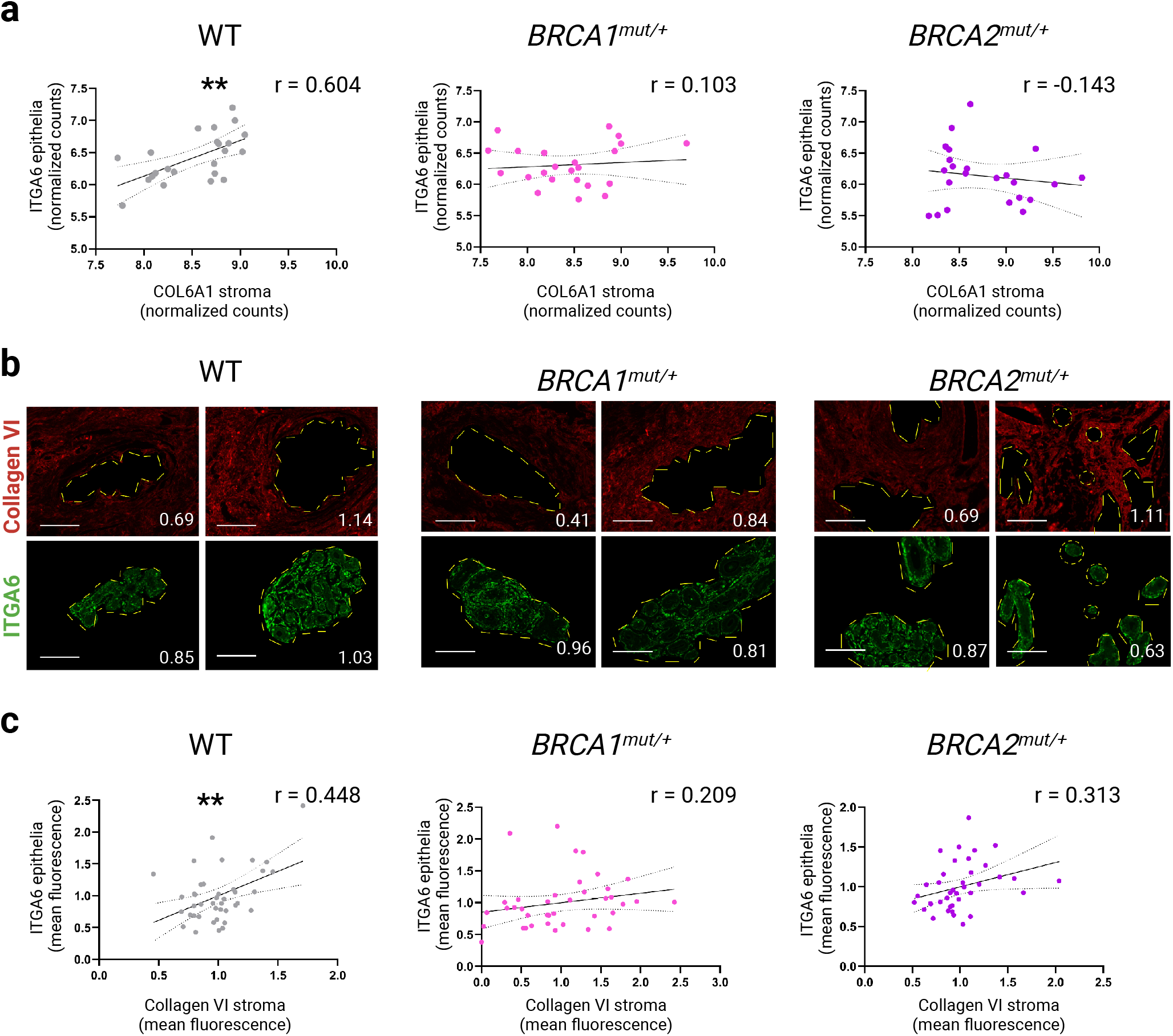
*ITGA6*-*COL6A1* expression is spatially correlated in non-mutation carriers but not in *BRCA1*/*2* mutation carriers. **a** *COL6A1* expression from stroma segments and *ITGA6* expression from epithelia segments are plotted on x and y axis, respectively. Each data point represents a single ROI, n = 23 for WT, *BRCA1*^*mut/+*^, and *BRCA2*^*mut/+*^ human breast tissues. The solid lines on the graph represent simple linear regression analysis, and the dashed lines represent 95% confidence intervals. The r value represents the Pearson correlation coefficient. **b** Representative images from immunofluorescence staining of consecutive FFPE tissue sections stained with type VI collagen (red) and ITGA6 (green) antibodies. Dashed yellow lines encircle the epithelial compartments within each field of view (FOV). Images show type VI collagen in the stroma only (epithelial region filled with black) and ITGA6 in the epithelia only (stromal region filled with black). Scale bar = 150 μm. Numbers at the bottom right represent normalized mean intensity of each image. **c** Quantification of images from the experiment performed in “b”. Results were collected from two independent experiments performed on a total of 12 patient (4 patients per genotype). Each data point represents a single FOV from WT (n = 41 images), *BRCA1*^*mut/+*^ (n = 40 images) and *BRCA2*^*mut/+*^ (n = 39 images). The statistical significance of Pearson correlations is calculated using an unpaired two-tailed t-test, ** p < 0.01.

## Discussion

Cell fate decisions are regulated by the microenvironment. Using whole-transcriptome digital spatial profiling of human breast tissues from control and *BRCA1/2* mutation carriers, we show for the first time that *BRCA1*^*mut/+*^ and *BRCA2*^*mut/+*^ preneoplastic epithelial cells communicate differently with their surrounding microenvironment. Our findings reveal both distinct and shared aspects of microenvironmental interactions in *BRCA1*^*mut/+*^ and *BRCA2*^*mut/+*^ breast tissues. We investigated autocrine and paracrine cell signaling through the identification of spatially defined receptor-ligand interactions. We found that integrin-promoted autocrine signaling in epithelial cells distinguished *BRCA2*^*mut/+*^ epithelial cells from *BRCA1*^*mut/+*^ epithelial cells. We discovered that expression of *ITGB1* and *ITGA11* correlated with the expression of type I collagen genes (*COL1A1, COL1A2*) in *BRCA2*^*mut/+*^ cells, but not in *BRCA1*^mut/+^ cells. Furthermore, we showed that in contrast to *BRCA1*^*mut/+*^ mammary epithelial cells, expression of multiple receptor-ligand pairs of TGF-*β*pathway, which is a major pathway required for collagen production, are differentially correlated in *BRCA2*^*mut/+*^ mammary epithelial cells. On the other hand, expression of *ITGB1* correlated with the expression of ECM degrading enzymes, growth factors or angiogenesis genes in *BRCA1*^*mut/+*^ epithelial cells compared to *BRCA2*^*mut/+*^ epithelial cells. The enhanced protease activity may cause subsequent ECM degradation that releases the ECM-bound growth factors and thereby increase their bioavailability in *BRCA1*^*mut/+*^ epithelial cells. In addition, we found that expression of collagen degrading enzymes *MMP3* and *MMP8* are increased in *BRCA1*^*mut/+*^ epithelial cells (Supplementary Figure 2a). Interestingly, a recent study discovered pre-cancer-associated fibroblasts that produce pro-tumorigenic MMP3, which promotes *BRCA1*-driven tumorigenesis in vivo [32]. Dysregulation of ECM can lead to several severe human conditions, including cancer. In some cases, the ECM becomes mechanically or enzymatically degraded. In other cases, a net accumulation of ECM is part of the pathologic event [33, 34]. Our data suggest that there is more collagen deposition in *BRCA2*^*mut/+*^ breast tissues, while there is more collagen degradation in *BRCA1*^*mut/+*^ breast tissues. This could explain why germline *BRCA1* and *BRCA2* mutations predispose to different subtypes of breast cancers. Regarding the paracrine signaling between epithelial cells and surrounding stromal cells, *BRCA1*^*mut/+*^ and *BRCA2*^*mut/+*^ preneoplastic breast tissues were comparable. In contrast to normal tissues, more integrin-ligand pairs were differentially correlated in *BRCA1/2*^*mut/+*^ breast tissues, with stromal cells expressing more integrin receptors. These findings suggest that communication from epithelia to surrounding stroma is increased in *BRCA1*/*2*^*mut/+*^ breast tissues. We hypothesize that epithelial cells may alter ECM composition in order to influence and educate stromal cells in *BRCA1*/*2* mutation carriers to acquire a pre-malignant phenotype.

Breast cancer is a heterogenous disease. Comprehensive gene expression profiling studies identified five major molecular subtypes: basal-like, luminal A, luminal B, HER2-positive and claudin-low [35, 36]. These subtypes are associated with different clinical outcomes and treatment responses [37, 38]. Our study investigated a critical question: why do germline *BRCA1* and *BRCA2* mutation carriers develop distinct subtypes of breast cancer? It is hypothesized that mammary stem cells and progenitors are the origin cells of breast cancer [39]. Both *BRCA1* and *BRCA2* mutation carriers demonstrate abnormal mammary epithelial cell differentiation [17-19]. Given the susceptibility of *BRCA1* and *BRCA2* mutation carriers to distinct subtypes of breast cancer, we suggest that *BRCA1* and *BRCA2* loss do not have the same effect on mammary epithelial cell differentiation. Clearly, microenvironmental cues play a crucial role in cell fate regulation and tissue-specific development. Here, we showed that in contrast to *BRCA1* deficiency, *BRCA2* deficiency increased the expression of activated *β*1 integrin and phosphorylated ERK in human mammary epithelial cells, reflecting an increase in force-mediated cellular mechano-signaling. High mechano-signaling is associated with the stiffness of the ECM and aggressiveness of the breast cancer. Furthermore, a stiff ECM may facilitate detrimental tumor characteristics by promoting stemness and a mesenchymal-like phenotype [40-42]. Amongst breast cancer subtypes, the invasive front of the most aggressive subtypes (basal-like and Her-2) was shown to be the stiffest and most heterogeneous, containing cells with the highest mechano-signaling, as compared to the less aggressive subtypes (luminal A and Luminal B) [43]. Importantly, it is well known that ECM is extensively remodeled at the advanced stages of cancer. However, less is known about the role of ECM in tumor initiation or in driving early phenotypic changes in high-risk breast tissues. Our findings provide evidence that *BRCA1* or *BRCA2* deficiency affects integrin-mediated mechano-signaling differently in normal human breast epithelial cells. Here, although we expanded on prior studies by dissecting epithelia-microenvironment communication in *BRCA1/2* mutation carriers, we were limited by the sample size. Nevertheless, we complemented our spatial transcriptomics-based findings with patient-derived cell culture models.

In conclusion, our work provides a framework for understanding the spatial interactions between mammary epithelial cells and the microenvironment directing epithelial cell fate in *BRCA1*/*2* mutation carriers and identifies markers for future translational studies aimed at improving risk prediction and breast cancer prevention in high-risk women.

## Methods

### Human breast tissues

Fixed human breast tissues were obtained from the Cleveland Clinic with approval by the local Institutional Review Board (protocol 19-1257). Details of patient characteristics are provided in Supplementary Table 1. The samples consisted of either normal breast tissues from reduction mammoplasties or noncancerous breast tissues from prophylactic mastectomies of known *BRCA1* or *BRCA2* mutation carriers. *BRCA1*/*2* mutation carrier status was determined by clinical germline genetic testing before tissue collection.

### Antibodies

Primary antibodies used for immunocytochemistry were 1:1000 KRT14 (BioLegend, 905301), 1:500 KRT18 (abcam, ab24561), 1:500 KRT8/18 (Invitrogen, MA5-14088), 1:300 active ITGβ1 (Sigma Aldrich, MAB2079Z), and 1:200 phospho-p44/42 MAPK (Erk1/2) (Thr202/Tyr204) (Cell Signaling Technology, 9101S). Primary antibodies used for immunohistochemistry of FFPE tissues were 1:50 COL6 (Rockland, 600-401-108-0.1) and 1:500 ITGA6 (abcam, ab181551). Secondary antibodies used for both methods, all diluted 1:1,000, were Alexa Fluor 488 Goat anti-Rabbit IgG Highly Cross-Adsorbed (Invitrogen, A-11034), Alexa Fluor 647 Goat anti-Mouse IgG Cross-Adsorbed (Invitrogen, A-21235), and Alexa Fluor™ 546 Goat anti-Rabbit IgG Cross-Adsorbed (Invitrogen, A-11010).

### GeoMx DSP profiling of the whole-transcriptome atlas (WTA)

Serially sectioned FFPE sections (5 μm) of specimens were prepared to generate consecutive sections that were processed for H&E and RNA hybridization. After drying overnight, the slides were baked at 42°C for 3.0 hours and again baked at 60°C for 3.0 hours just prior to deparaffinization. RNA assay slides were deparaffinized and subjected to antigen retrieval (Tris-EDTA, pH9.0; 15 minutes;>99°C), and enzymatic exposure of RNA targets (0.5μg/ml Proteinase K; 15 minutes; 37°C). Slides were then hybridized with human whole transcriptome overnight (nanoString, #GMX-RNA-NGS-HuWTA-4) at 37°C. The next morning slides underwent stringent washes and were prepared for morphology marker staining. Slides were treated with blocking buffer for 30 minutes at room temperature in a humidified chamber, then co-incubated with fluorescently labeled visualization markers; PanCK (pan-cytokeratin [CK], (Novus, Clone: AE1+AE3, Cat#: NBP2-34528), and with SYTO 13 (Thermo Fisher, Cat#S7575) for nuclei detection for 1 hour at room temperature in a humidified chamber. Slides were washed twice then loaded into the GeoMx for imaging. Once the samples were imaged, and regions of interest (ROIs) were selected, and any segmentation made. During collection, each capture area (whole ROI or segmented areas) was exposed to UV light that cleaved the linker and released the barcoded oligos for capture by microfluidics. The released barcodes were collected in 96-well plates and were used in the NGS readout library preparation procedure protocol. The resulted libraries were sequenced by the Illumina NextSeq 6000 platform using reversed, 2 × 27 base paired reads. The resulting GeoMx DSP data was then coupled to next generation sequencing (NGS) readout.

### Data preprocessing

Nanostring DSP quantification DCC files were analyzed with R version 4.1.3 [44]. R packages GeoMxWorkflows [45], GeomxTools [46], and NanoStringNCTools [47] were used to perform data normalization and quality control. Default values were used unless otherwise indicated. DCC counts and the PKC file (Whole Transcriptome Atlas, https://nanostring.com/wp-content/uploads/Hs_R_NGS_WTA_v1.0.pkc_.zip) were imported and segments identified as containing primarily ECM were excluded. All 0 counts were imputed to a value of 1. Candidate segment filter criteria were 1000 minimum reads, 80% trimming, 80% stitching, 80% alignment, 50% sequencing saturation, 10 reads as a minimum negative control count, 1000 reads as a maximum observed in NCT wells, and an area of 5000. Geometric means for negative and endogenous control probes were calculated. One segment was removed due to low negative and endogenous probe geometric means, which also had a sequencing saturation under 30%.

A global (across all segments) probe filter criteria was applied of a minimum probe ratio of 0.1 (geometric mean of a given probe’s counts across all segments, divided by geometric mean of all probes for the corresponding gene target across all segments) and if found to be an outlier via Grubb’s test in at least 20% of segments. Additionally, probes were filtered for a given segment if found to be an outlier according to Grubb’s test in that segment. Probe counts were aggregated to gene counts. A limit of quantification (LOQ) per segment was calculated as the product of the negative probes’ geometric mean and the square of negative probes’ standard deviation for each segment, or a value of 2, whichever was larger. Segments with less than 1% of genes above the LOQ were considered for exclusion. Genes were excluded if having a LOQ across segments < 1%, leaving 11,833 genes for downstream analysis. 75^th^ percentile (Q3) normalization was applied to the gene counts, which were then log2-transformed.

Visualization of segments before and after batch (scan) correction involved correcting log2 -normalized counts with ComBat [48], using scan as the batch indicator and genotype as the group indicator. Principal components were calculated for corrected and uncorrected counts and plotted with ggplot2 [49]. Plots of Q3 normalization and genes detected above LOQ were created with ggplot2.

### Differential gene expression analysis

Differential gene expression between genotype pairs was performed using linear mixed-effects models, which included genotype as a fixed effect, and a random intercept at the patient level. Testing was performed on ComBat-adjusted log2-normalized values within each segment (stroma and epithelium). The results were adjusted for multiplicity at an FDR threshold of 0.05 across each gene and genotype pairing. Genes with nominal significance (unadjusted p<0.05) were considered significant.

### Correlation Analysis

Known ligand-receptor pairs identified by Ramilowski 2015 [50] were downloaded from FANTOM (https://fantom.gsc.riken.jp/5/suppl/Ramilowski_et_al_2015/). Genes with average CAGE expression in the FANTOM mammary epithelial tissues were included for pairwise correlation analysis. Log2-normalized values were adjusted for batch by regressing out scan effect with linear regression and paired genes were compared between genotype pairs, both within each segment (Pearson correlation) and across segments (Spearman correlation) using a two-sized test. Multiple testing correction was performed with the Benjamini and Hochberg method, and adjusted p-values less than 0.05 were considered significant.

### Isolation and culture of HMECs from breast tissues

Breast tissues were collected and processed as previously described with minor modifications [19]. Briefly, the tissues were collected on ice, washed with PBS containing 50 U/mL penicillin-streptomycin, finely minced with scalpel, resuspended in 1-2 mg/mL collagenase solution (Sigma-Aldrich, C9722-50MG) in HBSS with Ca^2+^ and Mg^2+^ (Gibco, 24020) for one hour at 37 °C. The tissue was then dissociated into a single cell suspension, and filtered through 40 μm filter. Single cells were seeded onto collagen (Advanced BioMatrix, 5074) - coated plates in complete media: DMEM/F12 medium (ThermoFisher Scientific, 11330057) with 5% fetal bovine serum (Cleveland Clinic Foundation (CCF) Cell Culture Core), 5 μg/mL insulin (Sigma Aldrich, I1882-100MG), 2 mM L-glutamine (Gibco, 25030-081), 10 ng/mL human recombinant epidermal growth factor (PeproTech, AF-100-15), 500 ng/mL hydrocortisone (Sigma Aldrich, H0888-5G), 1x antibiotic/antimycotic (CCF Cell Culture Core) and 50 U/mL pen strep (CCF Cell Culture Core). Dead and floating cells were removed by replacing media daily. The cultures were initially obtained as mixed co-cultures of epithelial and stromal cells. Using differential trypsinization, the stroma was separated from the epithelia. Basal and luminal cytokeratin markers, KRT14 and KRT8/18, were co-immunostained on epithelial cultures to demonstrate the preservation of both basal and luminal mammary epithelial cell lineages.

siRNA transfections were carried out using Lipofectamine RNAiMAX (ThermoFisher Scientific, 13778075) according to the manufacturer’s protocol for “reverse transfection”. The siRNAs were provided as a “SMARTpool” of four separate siRNAs from Horizon Discovery (Supplementary Table 2).

### Real-Time qRT-PCR

Total RNA was isolated using TRIzol (Invitrogen, 15596026) according to manufacturer’s protocol. Complementary DNA synthesis was performed using PrimeScript RT Reagent Kit (Takara RR037A). The qPCR reactions were carried out using Power SYBR Green PCR Master Mix (Applied Biosystems 4367659) run on a 7500 Real Time PCR System (Applied Biosystems). *β-Actin* expression levels were used to normalize expression between samples and 2^ -ΔCT values were used for quantifying relative expression of genes, with siCTRL values set to a mean of 1. The primer sequences were listed on Supplementary Table 3.

### Immunocytochemistry and quantification

Cells were seeded on 8 well chamber slides (Ibidi, 80841) for immunofluorescence staining. Briefly, 50 μg/mL of Poly-D-Lysine (Sigma Aldrich P6407-5MG) in cell culture grade water (Invitrogen) was added to the wells of the chamber slides and allowed to incubate for one hour at room temperature to aid in cell attachment. This solution was then aspirated, followed by three washes with cell culture grade water. Slides were then allowed to dry at room temperature for one hour if being used immediately or wrapped in Parafilm (USA Scientific, 3023 -4526) and stored at 4 °C for later use. Slides were subsequently coated with collagen I solution as previously described in “Primary cell culture” section. Cells were then reverse transfected onto chamber slides, cultured for three days, and washed twice with PBS before being fixed in chilled 100% methanol for 10 minutes. After that, cells were washed twice with PBS containing 10% normal goat serum (Cell Signaling Technology, 5425S) before being permeabilized for 5 minutes with PBS containing 0.1% Triton X-100 (Sigma Aldrich, T8787-50ML). Cells were then washed and blocked for 1 hour at room temperature with PBS + 10% goat serum and 0.1% Triton X-100, then incubated with the appropriate primary antibodies overnight at 4 °C. Cells were then washed and incubated with fluorescently labeled secondary antibodies for 1 hour at room temperature. After washing, a drop of ProLong Diamond Antifade Mountant (Invitrogen, P36962) was added to each chamber and mounted with a glass coverslip (Electron Microscopy Sciences, 72200-40). Slides were then sealed with nail polish and imaged on EVOS M5000 microscope (ThermoFisher Scientific, A40486). Images (at least 10-20 fields of view (FOV) per siRNA condition) were analyzed using ImageJ software. The integrated density of the fluorescent channel corresponding to the protein of interest was measured after using the “color threshold” tool to set the same minimum brightness threshold to all samples. To quantify dual positive integrated density, the color threshold of the merged image was adjusted to filter out non-yellow regions within each FOV. These integrated density values were then divided by the number of nuclei in the same FOV, which was obtained by auto-thresholding the DAPI channel and applying the “watershed” and “analyze particles” tools in ImageJ, resulting in a mean fluorescence value for each FOV.

### Immunohistochemistry and quantification

Slides of sectioned FFPE patient tissues were baked at 62 °C for 1 hour, deparaffinized with three washes in xylenes for five minutes each, and rehydrated in sequentially decreasing concentrations of ethanol for ten minutes each. Antigen retrieval was performed in 10mM sodium citrate buffer pH 6.0 (for COL6) or 10 mM Tris / 1 mM EDTA buffer pH 9.0 (for ITGA6) at 110 °C for 17 minutes in a Decloaking Chamber NxGen from BioCare Medical. After cooling to room temperature, slides were blocked in PBS + 10% goat serum for 1 hour and incubated with primary antbodies overnight at 4 °C in PBS + 0.5% BSA. Cells were then washed three times with PBS and incubated with secondary antibodies for one hour at room temperature in PBS + 0.5% BSA. After 4 washes with PBS, slides were mounted with ProLong Diamond Antifade Mountant, coverslipped, and sealed with nail polish. Imaging occurred on an EVOS M5000. Images were then analyzed with ImageJ software. To quantify epithelial ITGA6 or stromal COL6, epithelial regions were outlined based on morphology of DAPI images. The selected region was then transferred to the channel of the protein of interest, and either the “clear outside” or “fill” tool was applied to block out the stromal or epithelial regions, respectively. “Color threshold” was then applied to set the same minimum brightness to all samples, and the mean intensity value was recorded. After subtracting the mean background obtained from no primary antibody control slides using this same method, mean intensity values for ITGA6 and COL6 were normalized separately by dividing each value by the average of its genotype within an experiment.

## Data availability

The raw and processed RNA-sequencing data generated in this study will be available in Gene Expression Omnibus (GEO). The remaining data are available within the Article and Supplementary Information.

## Code availability

Code for gene expression quality control and downstream analyses will be made available on github.

## Acknowledgements

We thank the Advanced Genomics Core at the University of Michigan for library preparation and next-generation sequencing. We are very grateful to Cleveland Clinic Central Biorepository for providing us human breast tissues. We also thank pathologists Drs. Xiaoyan Cui and John Van Arnam for their help with patient sample access. All of the figures were created by Biorender. This work was supported by seed funds from the Cleveland Clinic’s Lerner Research Institute (M.K.), R01CA213843 (R.A.K.), R01CA257502 (R.A.K.), and VeloSano for the Cure (R.A.K.).

## Author contributions

M.K. designed and supervised the project. E.D.K. and P.M. contributed patient samples. E.D.K. prepared tissue slides for spatial transcriptomics. M.K., A.C and K.V. designed and performed the experiments, and interpreted data. P.B. performed the bioinformatic analyses with help from B.H., Y.N. and A.C. M.K and A.C. wrote the manuscript with scientific insights from R.A.K. A.C., K.V. and P.B. generated the figures. All authors read and contributed to editing the manuscript.

