## Supplementary Information for "Alterations in the preneoplastic breast microenvironment of *BRCA1/2* mutation carriers revealed by spatial transcriptomics"

**Supplementary Figure 1: Digital spatial profiling of human breast tissues.**

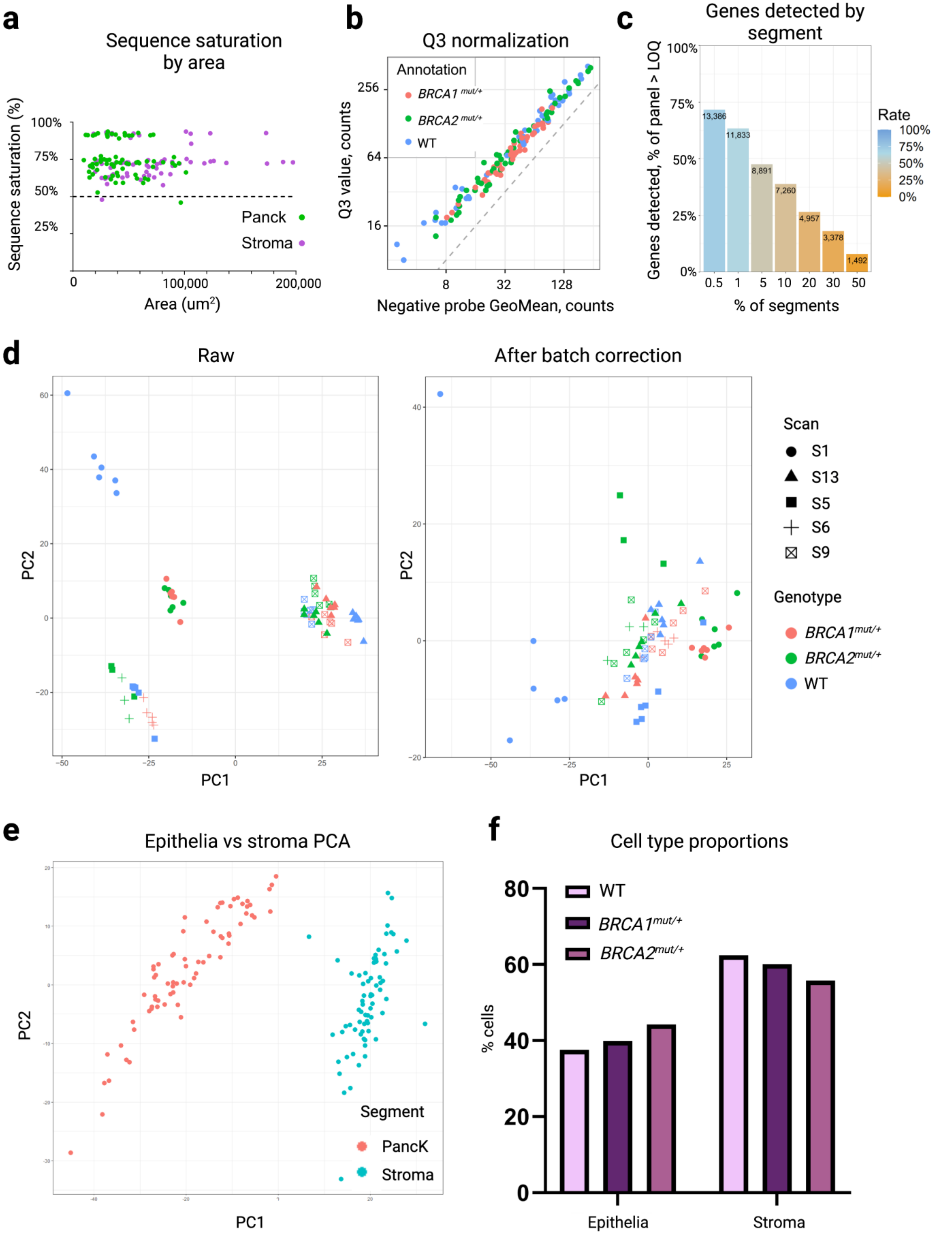

**g**

**Epithelia**

| WT - *BRCA1*$^{mut/+}$ | p-value |
|---|---|
| 1. LDLRAD4 and what we know about it | 2.12E-03 |
| 2. Heme Biosynthesis | 4.78E-03 |
| 3. Interleukin-1 Induced Activation of NF-kappa-B | 6.62E-03 |
| 4. Metabolic reprogramming in colon cancer | 6.90E-03 |
| 5. Rett syndrome causing genes | 1.31E-02 |
| 6. Endoderm differentiation | 1.32E-02 |
| 7. G1 to S cell cycle control | 1.54E-02 |
| 8. ESC Pluripotency Pathways | 2.08E-02 |
| 9. Osteoblast differentiation | 2.31E-02 |
| 10. Wnt signaling pathway and pluripotency | 2.40E-02 |

| WT - *BRCA2*$^{mut/+}$ | p-value |
|---|---|
| 1. Apoptosis-related network due to altered Notch3 in ovarian cancer | 3.89E-05 |
| 2. Photodynamic therapy-induced unfolded protein response | 4.31E-04 |
| 3. Brain-derived neurotrophic factor (BDNF) signaling pathway | 8.01E-04 |
| 4. 4-hydroxytamoxifen, Dexamethasone, and Retinoic Acids Regulation of p27 Expression | 8.04E-04 |
| 5. Type II diabetes mellitus | 1.78E-03 |
| 6. Insulin Signaling | 1.88E-03 |
| 7. Head and Neck Squamous Cell Carcinoma | 1.96E-03 |
| 8. EGF/EGFR signaling pathway | 2.07E-03 |
| 9. IL-1 signaling pathway | 2.11E-03 |
| 10. Unfolded protein response | 2.49E-03 |

**Stroma**

| WT - *BRCA1*$^{mut/+}$ | p-value |
|---|---|
| 1. Arrhythmogenic Right Ventricular Cardiomyopathy | 9.40E-06 |
| 2. Focal Adhesion | 2.12E-05 |
| 3. Focal Adhesion-PI3K-Akt-mTOR-signaling pathway | 4.01E-05 |
| 4. MFAP5 effect on permeability and motility of endothelial cells via cytoskeleton rearrangement | 4.86E-05 |
| 5. Primary focal segmental glomerulosclerosis (FSGS) | 4.95E-05 |
| 6. Thyroid hormones production and their peripheral downstream signaling effects | 8.50E-05 |
| 7. VEGFA-VEGFR2 Signaling Pathway | 1.11E-04 |
| 8. Brain-derived neurotrophic factor (BDNF) signaling pathway | 1.68E-04 |
| 9. PI3K-Akt signaling pathway | 1.93E-04 |
| 10. Prolactin Signaling Pathway | 4.32E-04 |

| WT - *BRCA2*$^{mut/+}$ | p-value |
|---|---|
| 1. TCA Cycle (aka Krebs or citric acid cycle) | 3.54E-03 |
| 2. Proteoglycan biosynthesis | 3.54E-03 |
| 3. Translation Factors | 2.09E-03 |
| 4. Canonical and non-canonical Notch signaling | 1.13E-02 |
| 5. Intraflagellar transport proteins binding to dynein | 1.13E-02 |
| 6. Benzene metabolism WP3891 | 1.00E-01 |
| 7. NOTCH1 regulation of endothelial cell calcification | 3.47E-02 |
| 8. Aflatoxin B1 metabolism | 1.16E-01 |
| 9. Disorders of the Krebs cycle | 1.16E-01 |
| 10. MicroRNA for Targeting Cancer Growth and Vascularization in Glioblastoma WP3593 | 1.16E-01 |

**a** Except for two ROIs, RNA-sequencing saturation graph showing greater than 50% saturation for all segments. **b** Q3 normalization reveals that all endogenous probe counts for all ROIs are greater than the negative control probes. **c** Genes detected across percent of segments. The 11,833 genes present in at least 1% of segments were used as the cutoff for downstream analysis. **d** Principal component analysis (PCA) annotated by genotype and batch (scan), demonstrating a significant batch effect in raw data (left) vs. after ComBat batch correction (right). **e** PCA of all ROIs showing separation of epithelial and stromal segments after batch correction. **f** Analysis of area occupied by epithelial and stromal cell types across ROIs. Bars represent mean values. **g** Top-ranked WIKI pathway analysis results of *WT-BRCA1*$^{mut/+}$ (left) and *WT-BRCA2*$^{mut/+}$ (right) DEGs in epithelia (top) and stroma (bottom) from i and ii in Fig. 1b.

**Supplementary Figure 2: Spatial analysis of epithelial and stromal cells reveals altered ECM signaling in *BRCA1* and *BRCA2* mutation carriers.**

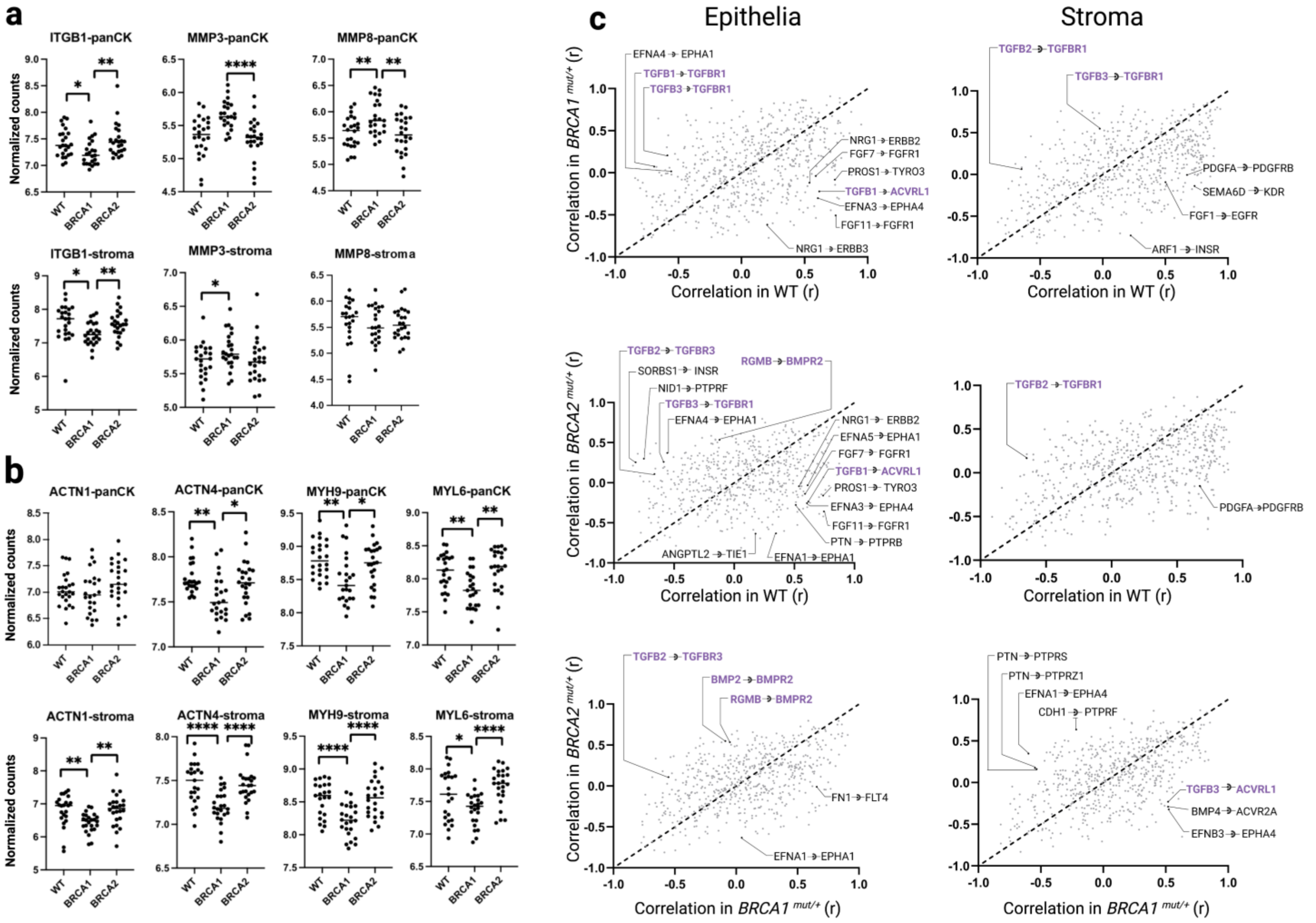


Batch-corrected normalized counts of ROIs from each genotype comparing epithelial (top) and stromal (bottom) gene expression for **a** *ITGB1*, matrix metalloproteinases *MMP3* and *MMP8*, and **b** actin/myosin genes. Statistical significance was determined using unpaired two-tailed t-tests with * $p < 0.05$, ** $p < 0.01$ and **** $p < 0.0001$. **c** Pearson correlation coefficients of wild-type (WT), *BRCA1*$^{mut/+}$ and *BRCA2*$^{mut/+}$ patients plotted within epithelial (left) and stromal (right) compartments. Differentially correlated enzyme-linked receptor-ligand pairs are labeled. TGF$\beta$ receptors-cognate ligand pairings are highlighted in purple.

**Supplementary Figure 3: Human breast epithelial cell cultures.**

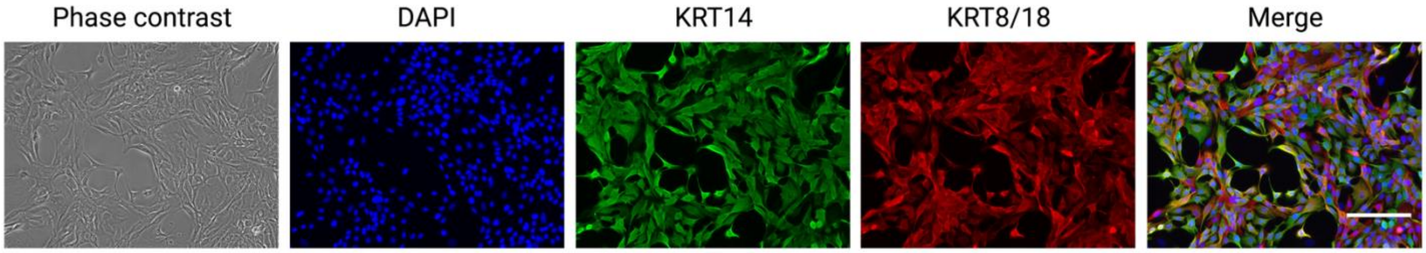


Primary human breast epithelial cell cultures were immuno-stained for luminal epithelial (KRT8/18) and basal epithelial (KRT14) cyto-keratins to demonstrate the preservation of both epithelial lineages under our culture conditions.

**Supplementary Figure 4: Expression of spatially correlated receptor-ligand pairs.**

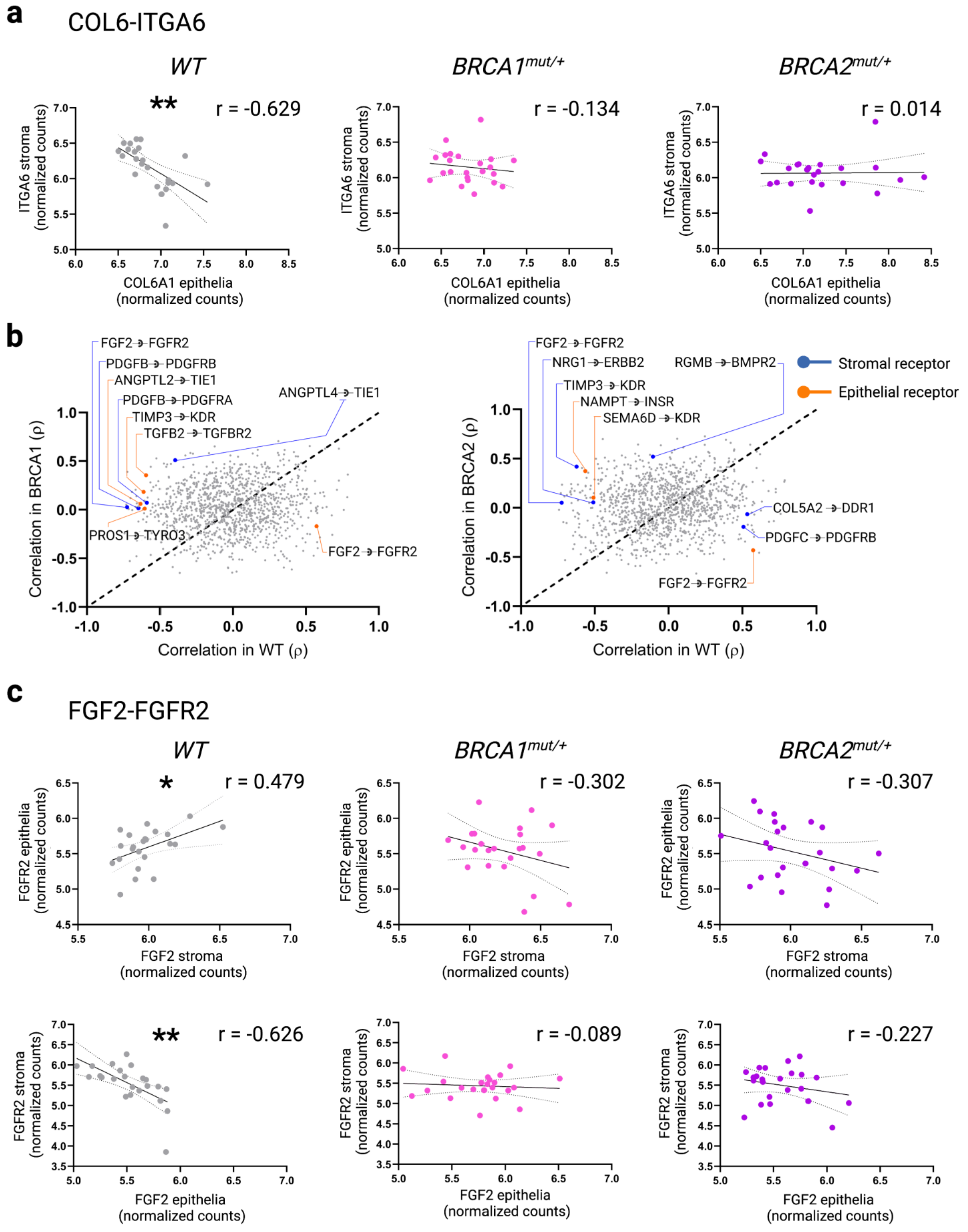

**a** COL6A1 expression from epithelia and ITGA6 expression from stroma segments are plotted on x and y axis, respectively. **b** Enzyme-linked receptor-ligand pairs that were differentially correlated in pairwise comparison across genotypes are labeled. Blue dots and lines indicate a stromal receptor. Orange dots and lines indicate an epithelial receptor. **c** FGFR2-FGF2 expression is spatially correlated in non-mutation carriers but not in *BRCA1/2* mutation carriers.

### Supplementary Table 1: Patient characteristics

| Patient ID | Age | Genotype | Variant | Parity | Hormonal status | Mammographic Density |
|---|---|---|---|---|---|---|
| BRCA1-1 | 28 | *BRCA1* $^{mut/+}$ | 5296del4 aka [c.5177_5180del (p.Arg1726Lysfs*3)] | G3P0 | premenopausal | focal enhancement |
| BRCA1-2 | 43 | *BRCA1* $^{mut/+}$ | W 1508X (4643 G>A) | G3P3 | premenopausal | focal enhancement |
| BRCA1-4 | 44 | *BRCA1* $^{mut/+}$ | c.4964_4982del | G1P1001 | Postmenopausal | Negative |
| BRCA1-6 | 40 | *BRCA1* $^{mut/+}$ | c.676del | G3P1021 | Postmenopausal | Negative |
| BRCA2-1 | 30 | *BRCA2* $^{mut/+}$ | c.9182T>A (p.Leu3061*) | G1P1 | premenopausal | Negative |
| BRCA2-2 | 45 | *BRCA2* $^{mut/+}$ | c.3922G>T (p.Glu1308*) | G3P3 | postmenopausal | R focal enhancing mass |
| BRCA2-4 | 42 | *BRCA2* $^{mut/+}$ | c.7069_7070del (p.Leu2357Valfs*2). | G5P2 | premenopausal | Negative |
| BRCA2-6 | 40 | *BRCA2* $^{mut/+}$ | c.3708dup | G2P2002 | premenopausal | diffuse enhancement |
| WT-1 | 25 | WT | N/A | G1P1001 | premenopausal | Negative |
| WT-2 | 40 | WT | N/A | G0P0000 | premenopausal | Negative |
| WT-4 | 42 | WT | N/A | NA | premenopausal | Negative |
| WT-6 | 39 | WT | N/A | G4P3104 | premenopausal | Negative |

### Supplementary Table 2: siRNA targeted sequences

| siRNA | Targeted sequences |
|---|---|
| siCTRL | UGGUUUACAUGUCGACUAA, UGGUUUACAUGUUGUGUGA, UGGUUUACAUGUUUUCUGA, UGGUUUACAUGUUUUCCUA |
| siBRCA1 | CAACAUGCCCACAGAUCAA, CCAAAGCGAGCAAGAGAAU, UGAUAAAGCUCCAGCAGGA, GAAGGAGCUUUCAUCAUUC |
| siBRCA2 | GAAACGGACUUGCUAUUUA, GGUAUCAGAUGCUUCAUUA, GAAGAAUGCAGGUUUAAUA, UAAGGAACGUCAAGAGAUA |

### Supplementary Table 3: qRT-PCR primer sequences

| Gene | Forward primer | Reverse primer |
|---|---|---|
| *β-Actin* | CATGGATGATGATATCGCCGC | CACGATGGAGGGGAAGACG |
| *BRCA1* | GTCCCATCTGTCTGGAGTTGA | GGCCCTTTCTTCTGGTTGAGA |
| *BRCA2* | CCTGATGCCTGTACACCTCTT | GCAGGCCGAGTACTGTTAGC |
